# Emotional content effects the precision of visual working memory

**DOI:** 10.1101/2024.12.01.626217

**Authors:** Zeinab Haghian, Abdol-Hossein Vahabie, Ehsan Rezayat

## Abstract

**Objectives:** Our working memory (WM) is susceptible to errors influenced by various sources. Recent research has illuminated the intricate relationship between emotional valence and working memory performance. This study aims to comprehensively investigate memory recall biases across different emotional categories.

**Method:** To explore this relationship, we designed and implemented a delayed-reproduction facial emotion n-back task. Participants were tasked with recalling the emotional valence of target faces across various n-back conditions, selecting their responses from a continuum of 19 morphed faces ranging from sad to happy.

**Results:** Our findings indicate that participants generally exhibit a negativity bias, struggling more with happy faces. Interestingly, they also perceive happy faces as less happy and sad faces as less sad, suggesting both positive and negative reappraisal in their emotional valence perception. This underscores the complex interplay between emotional valence and cognitive performance. Notably, recall of neutral images remained stable and was unaffected by preceding emotional contexts.

**Conclusion:** These findings demonstrate that emotional content in working memory significantly impacts errors during WM tasks, with more emotionally charged faces leading to a greater drift toward the lower valence axis. This highlights the need for further exploration of how emotional factors influence cognitive processes in working memory.

## 1. Introduction

Working memory (WM) is essential for goal-directed behaviors, facilitating the temporary storage and manipulation of information necessary for complex cognitive tasks such as reasoning, learning, problem-solving, and comprehension (Chai et al., 2018; Diamond, 2013; Malenka et al., 2009; Miyake & Shah, 1999). As a central component of cognition, WM interacts intricately with other cognitive processes, notably attention, long-term memory, and emotion. Among these, the relationship between WM and emotion has garnered significant interest due to its profound and bidirectional effects. Emotional experiences can enhance or impair WM performance, influenced by factors such as the intensity and valence of the emotion, as well as the cognitive demands of the task (Mikels & Reuter-Lorenz, 2019; Hou et al., 2022). Conversely, WM is crucial for managing and manipulating emotional information, supporting emotional regulation and adaptive decision-making. Thus, the interplay between WM and emotion is complex and context-dependent, reflecting their mutual influence on cognitive and affective functioning.

The role of emotion in WM has significant implications for human behavior, particularly in detecting and responding to survival-relevant situations. Emotional stimuli, especially facial expressions, serve as vital social cues and are among the most studied in this context. Research shows that emotional faces are processed more efficiently than neutral ones, highlighting the evolutionary importance of swiftly interpreting emotional signals (Johansson et al., 2004). The emotional significance of a stimulus often enhances its processing efficiency, allowing for quicker detection and evaluation (Zeelenberg et al., 2006). Emotional stimuli can capture cognitive resources more effectively than neutral stimuli, potentially facilitating or disrupting WM performance depending on the context. For instance, high-intensity emotions can demand considerable cognitive resources, possibly impairing WM efficiency (Hou et al., 2022). Despite advancements in understanding these interactions, debates persist regarding the differing impacts of positive and negative emotions on WM performance. Some studies suggest that positive emotions broaden cognitive scope, aiding WM (Fredrickson, 2001), while others argue that negative emotions, like sadness or anxiety, may narrow attentional focus, enhancing task-specific performance (Gasper & Clore, 2002; Shen et al., 2023; Kuehne et al., 2021).

A key factor in emotional processing is valence, which denotes the intrinsic positivity or negativity of a stimulus. Researchers have explored whether stimuli at the extremes of the valence continuum—positive (e.g., happy) or negative (e.g., sad or angry)—receive preferential processing. This has led to the identification of emotional processing biases, such as positive bias (favoring positive stimuli) and negative bias (favoring negative stimuli) (Dolcos et al., 2021; Xu et al., 2021). While evidence supports both biases, their nature and mechanisms remain actively investigated. Negative biases are often linked to the evolutionary significance of threat detection, while positive biases may arise from the frequent occurrence of positive stimuli in daily life (Esteves & Öhman, 1993; Johnston et al., 2001).

To study WM, researchers often use paradigms that require participants to determine if a presented object matches a previously seen one, yielding binary outcomes (correct or incorrect) (Bays et al., 2009). However, these methods can limit the assessment of memory retention quality. Alternative paradigms, such as “delayed-reproduction” tasks, offer continuous measures of WM performance, allowing for a more nuanced understanding of how items are retained and manipulated (Gorgoraptis et al., 2011). Our study employed a delayed-reproduction facial emotion N-Back task using morphed emotional faces to investigate the interaction between emotional content and WM performance. This design enabled precise measurement of perceptual errors and reaction times, facilitating the exploration of positive and negative biases in recalling emotional stimuli. By examining how emotional valence and intensity influence cognitive processes, our research contributes to understanding emotion-cognition interactions, with implications for emotional regulation, adaptive behavior, and everyday functioning.

## 2. Materials and Methods

### 2.1 Participants

A total of 31 healthy undergraduates and graduate students, (17 female, *age*: [19, 38], mean ± *SD*: 24.3 ± 3.05) volunteered to attend this experiment after providing written informed consent online to procedures approved by the local ethics committee. All participants had normal or corrected-to-normal visual acuity. All procedures were approved by University of Tehran Science Committee (**Approval ID IR.UT.PSYEDU.REC.1403.012**) and was in accordance with Helsinki declaration of 1964 and its revisions.

### 2.2 Stimuli and Apparatus

We created a neutral face (emotion and sex-neutral) using FaceGen Modeller 3.5 (Singular Inversions, ON, Canada). This face was then morphed into two categories, sad and happy, at 10% intervals using InterFace software (Kramer et al., 2017). This process resulted in 19 morphed face stimuli, ranging from 90% happy to 90% sad (Figure 1A). All morphed faces were then cropped into oval shapes, using Adobe Photoshop software and presented on a screen with a gray background. Luminance was controlled for in the emotional faces using the SHINE toolbox in MATLAB (Willenbockel et al., 2010). We considered a series of images, with Image 1 depicting 90% sadness and Image 19 depicting 90% happiness, while the intermediate images were arranged sequentially. In our study, images 1 through 7 were categorized as sad, images 8 through 12 were categorized as neutral, and images 13 through 19 were categorized as happy. Face stimuli (3.33°× 4.76°) were presented on the center of a 23-inch TFT monitor with a resolution of 1920 × 1080 pixels and a refresh rate of 60 Hz. The experiment was programmed using the PsychToolbox 3 extension (Brainard, 1997) running in MATLAB (2019b) (MathWorks, Natick, MA, United States).

**FIGURE 1.**
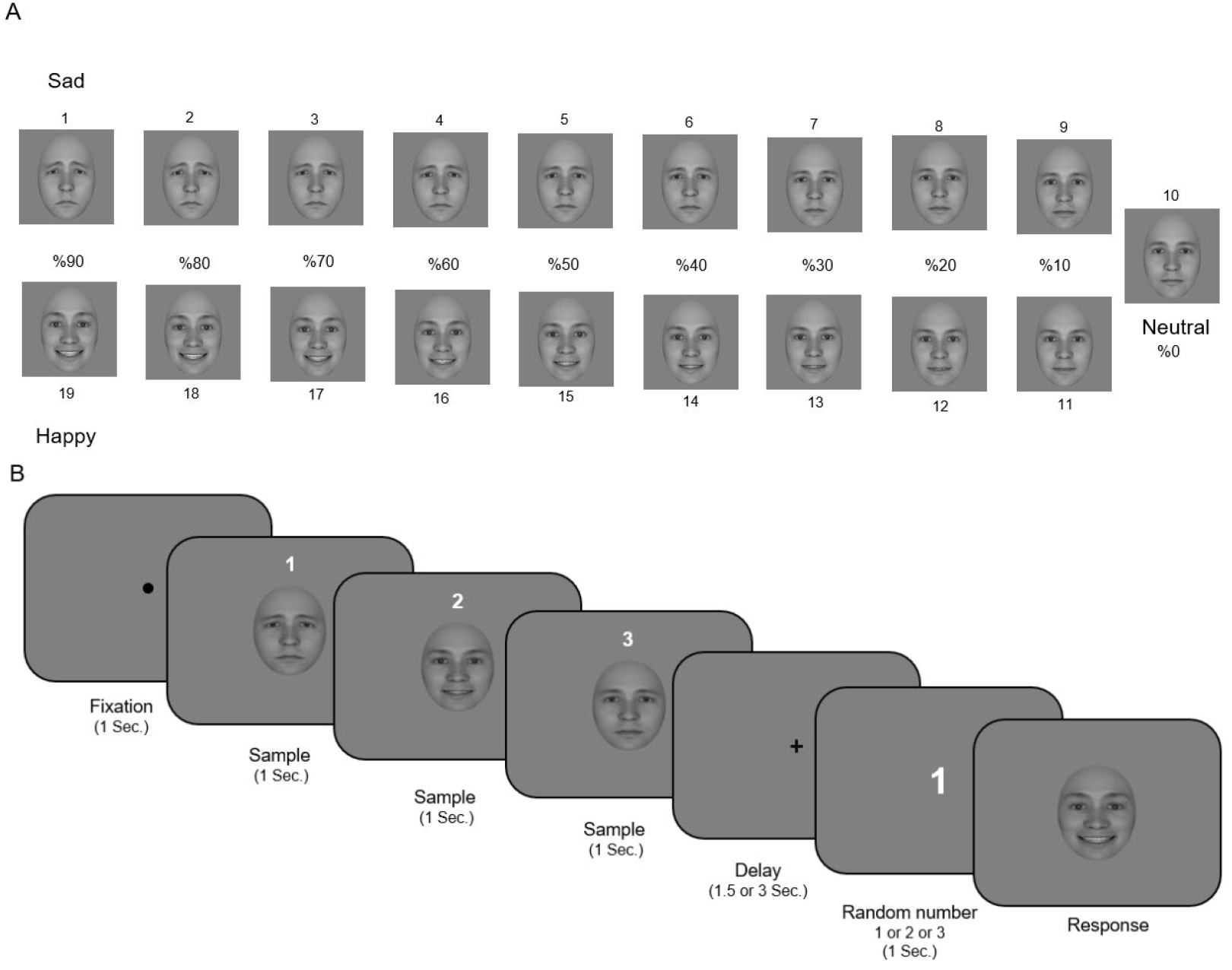
Experiment procedure and task. (A) Continuum of Morphed Facial Stimuli. A series of facial stimuli created through a morphing process. (B) The schematic description of the trial sequence for behavioral task. Each trial began with presenting a fixation dot for 1 second, followed by three ‘sample face’ presentation (1 second for each one) and a delay (1.5 or 3 seconds). A number were appeared on screen to show which image would be remembered. Then A ‘test face’ was presented and the subject had to change the stimulus to match it to the remembered ‘sample face’.

### 2.3 Procedure

Participants sat comfortably on a chair at a fixed distance of ∼60 cm from the screen in a dimly lit room, with their heads resting on a chinrest. Before the main task, instructions were displayed on the screen, and the examiner explained the procedure to each participant. Participants completed five practice trials to familiarize themselves with the task. Each trial began with a fixation spot (black, 0.3° diameter, Figure 1B), on which participants were instructed to maintain their gaze. After 1 second, three face stimuli (“**sample faces**”) appeared on the screen simultaneously, each for one second, with numbers 1, 2, or 3 displayed above the respective faces. These faces were selected pseudorandomly from 0%, 20%, 40%, 60%, or 80% morphed faces, with one of the three faces serving as the target face.

Following the presentation of the sample faces, a fixation cross (black, 0.2° length of each line) appeared at the center of the screen for a delay period of either 1.5 or 3 seconds, chosen pseudorandomly for each trial. Participants were instructed to maintain their gaze on the fixation cross during this delay. After the delay, a number (1, 2, or 3) corresponding to one of the sample faces was displayed on the screen for one second. Subsequently, a random face (“**response face**”), which could be any of the 19 morphed faces, was presented. Participants were instructed to use the “Up” and “Down” arrow keys on the keyboard to adjust the “response face” through the morphs to match it to the “target face” corresponding to the displayed number (1, 2, or 3). They pressed the “ENTER” key to submit their responses. Each block consisted of 54 trials (2 delays × 9 sample faces × 3 repetitions). Each participant completed three blocks, with the option to take a 3–5 minutes break between blocks. We recorded participants’ choices and reaction times for each trial. No feedback was provided regarding their responses or reaction times.

### 2.4 Data analysis

To quantify participants’ perception accuracy, we defined the error on each trial as the difference between the target face and the participant’s selected face, calculated as (E*rror* = *Response* - *Target*). This error ranged from -17 to +17, where positive errors indicated that participants perceived the image as more pleasant in valence than it was, and negative errors indicated that they perceived it as more unpleasant in valence. For instance, if the target face was Image 16 (60% happy) and the participant selected Image 13 (30% happy), the error would be -3, reflecting a perception of less happiness than it actually was. This method allowed us to systematically assess the accuracy of emotional recognition in our participants.

We applied the generalized linear mixed-effects model (GLMM) for this study, which is specified to analyze the effects of reaction time (RT), intensity, and cognitive load (N-Back) on error rates across trials and participants for each emotion (Sad, Neutral, Happy). The model is expressed as:

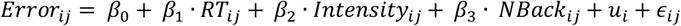

where *Error*_*ij*_ represents the error for participant *i* on trial *j, β*_0_ is the fixed-effect intercept, and *β*_1_, *β*_2_, and *β*_3_ are fixed-effect coefficients quantifying the influence of RT, intensity, and N-Back, respectively. The term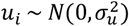 captures participant-specific variability as a random intercept, while *ϵ*_*ij*_∼ *N*(0, σ^2^) represents residual trial-level error. This GLMM structure enables the simultaneous assessment of both fixed effects, which are consistent across participants, and random effects, which account for individual differences. By incorporating hierarchical variance components, the model provides a robust framework for examining the predictors of error in perceptual accuracy while controlling for within-subject dependencies (Bolker et al., 2009; Bates et al., 2015).

## 3. Results

Our findings show that participants recalling happy images indeed tended to have lower precision (higher mean absolute errors) across all n-back conditions compared to those recalling sad or neutral images (Figure 2A). Participants also have less precision (made significantly more errors) in the happy condition compared to the sad condition (p = 0.0201). This finding suggests that the emotional state of happiness may impair cognitive performance, leading to a higher error rate. However, there were no significant differences in errors between the sad and neutral conditions (p = 0.7124), nor between the neutral and happy conditions (p = 0.1531). A one-way ANOVA was conducted to examine the effect of emotional categories (Sad, Neutral, Happy) on response errors. The results indicated a significant effect of emotion on errors at the (p < 0.05) level for the three conditions [F(2, 5019) = 324.48, p < 0.001] (Figure 3). Post hoc comparisons using the Bonferroni correction revealed significant differences between the groups: Sad vs. Neutral (mean difference = 1.3483, (p < 0.0001)), Sad vs. Happy (mean difference = 2.6661, (p < 0.0001)), and Neutral vs. Happy (mean difference = 1.3178, (p < 0.0001)). Moreover, the reaction times also have similar results (Figure 2B), participants had significantly longer RTs in the neutral condition compared to the sad condition (p = 0.0413). This suggests that the neutral emotional state may slow down cognitive processing speed compared to the sad state. Sadness might heighten focus and vigilance, leading to faster reaction times, whereas a neutral state might lack the emotional arousal needed to enhance cognitive performance. Additionally, participants had significantly longer RTs in the happy condition compared to the sad condition (p = 0.0008), indicating that happiness may impair cognitive performance by slowing down reaction times. There were no significant differences in RTs between the neutral and happy conditions (p = 0.1770), suggesting that both emotional states had a similar impact on cognitive processing speed. Despite these observed patterns, the ANOVA results showed no significant differences in errors or reaction times (RTs) across the different n-back conditions (1-back, 2-back, 3-back). This suggests that the cognitive load imposed by the varying n-back tasks did not significantly affect participants’ performance in terms of errors or RTs. Participants appeared equally adept at handling the cognitive demands of each n-back condition, indicating that the differences in cognitive load were not substantial enough to produce measurable effects in this context.

**Figure 2.**
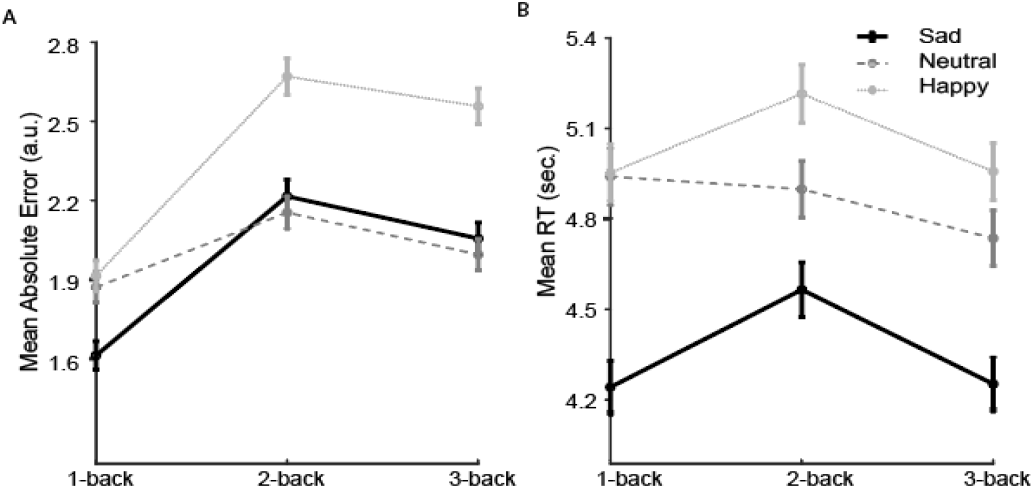
Happy content have more Mean Errors and Reaction Times for Emotional Recognition Across N-back Conditions. (A) Mean absolute errors for each emotion (sad, neutral, happy) across three n-back conditions (1-back, 2-back, 3-back). (B) Mean reaction times (RTs) for the same emotional categories and n-back conditions error bar is standard error across subjects.

**FIGURE 3.**
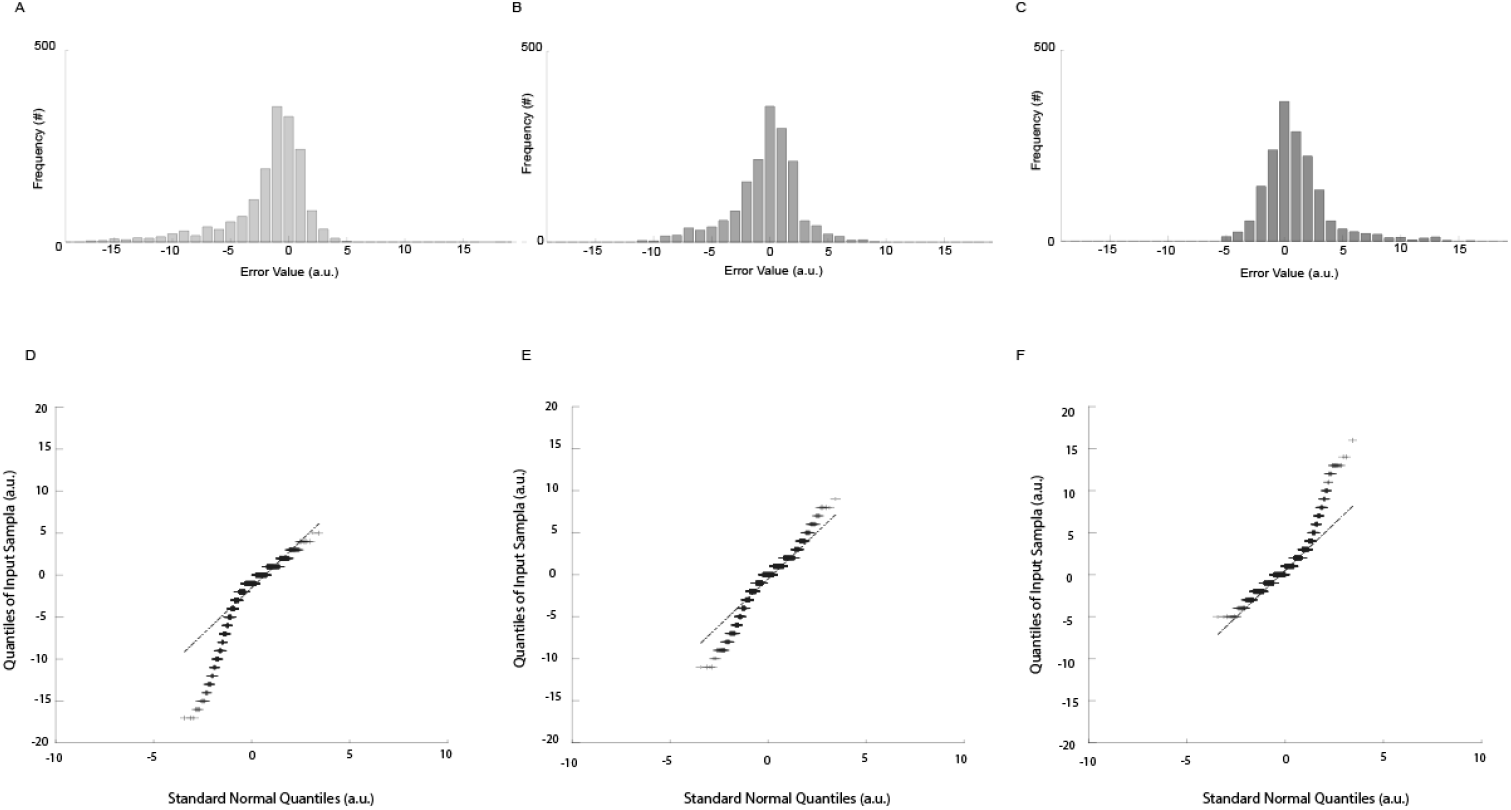
Error Distributions for Emotional Recognition were shifted in opposite direction of valance axis. The x-axis quantifies the error value, reflecting the deviation of the participant’s response from the target image. The y-axis enumerates the frequency of occurrences for each error value, corresponding to the number of images selected by participants. (A) represents the distribution for happy emotions, (B) for neutral emotions, and (C) for sad emotions. D-F, Quantile-Quantile Distributions for Emotional Recognition Errors. (D) represents the distribution for happy emotions, (E) for neutral emotions, and (F) for sad emotions.

We also found that participants recalled happy images as more unpleasant valence and sad images as more pleasant valence than they were initially presented. The analysis of skewness for the error distributions across different emotional categories reveals significant insights into the asymmetry of the data (Figure 3). The error distribution for the Happy category exhibits a negative skewness (-1.7198), indicating a higher frequency of extreme negative errors. This suggests that when participants attempt to recall happy images, they tend to make larger negative errors, meaning they chose images with a more unpleasant valence compared to the response image. In contrast, the Neutral category shows a slight negative skewness (-0.7085), reflecting a relatively balanced distribution of errors with a minor tendency towards larger negative errors. This indicates a slight preference for sadder images over happier ones in valence when recalling neutral images. The Sad category, however, displays a positive skewness (1.5368), indicating a higher frequency of extreme positive errors. This implies that when participants attempt to recall sad images, they are more prone to making larger positive errors, meaning they chose images with a more pleasant valence compared to the response image.

Using GLMM for the emotions of Sad, Neutral, and Happy, we also investigate the relationship between the predictor variables (Reaction Time (RT), Intensity, and N-Back) and the dependent variable (Error). The results of the regression analyses for the sad, neutral, and happy emotional categories reveal distinct relationships between error rates and predictors, including response time (RT), intensity, and cognitive load (N-Back). Across all three emotional categories, RT and intensity emerged as significant predictors of error rate, although the direction and magnitude of these relationships varied. The N-Back predictor, representing cognitive load, generally did not reach significance, suggesting it has a limited direct impact on error rates when controlling for RT and intensity.

**Table 1.** Generalized Linear Mixed Model Results.

| *Emotion* | *Predictor* | *Coefficient* | *p-value* | *Marginal*<br>*R<sup>2</sup>* | *Conditional*<br>*R<sup>2</sup>* |  |
| --- | --- | --- | --- | --- | --- | --- |
| *Sad* | Intercept | 1.8944 | <0.001 | 0.1107 | 0.1750 |  |
|  | RT | 0.12106 | <0.001 |  |  | Faster reaction times are associated with increased error |
|  | Intensity | 0.38182 | <0.001 |  |  | Higher intensities are associated with more positive error (more pleasant) |
|  | N-Back | 0.026362 | 0.75829 |  |  |  |
| *Neutral* | Intercept | -0.56018 | 0.25513 | 0.0364 | 0.0546 |  |
|  | RT | 0.16644 | <0.001 |  |  | Faster reaction times are associated with increased error |
|  | Intensity | -0.03816 | 0.35809 |  |  |  |
|  | N-Back | -0.11629 | 0.19824 |  |  |  |
| *Happy* | Intercept | 3.1251 | <0.001 | 0.1507 | 0.2750 |  |
|  | RT | 0.098246 | 0.00561 |  |  | Faster reaction times are associated with increased error. |
|  | Intensity | -0.32623 | <0.001 |  |  | Higher intensities are associated with more negative error (less pleasant) |
|  | N-Back | -0.050397 | 0.61385 |  |  |  |

Table 2, shows the Generalized Linear Mixed Model Results, which indicate the explanatory power of the fixed effects and the full model, respectively. For Sad and Happy emotions, the models show higher explanatory power, with marginal *R*^2^ values. This demonstrates that both the fixed predictors (RT, Intensity, and N-Back) and random effects (subject variability) contribute meaningfully to explaining Error. In these emotions, Intensity emerges as a significant predictor, with higher intensities reducing Error. Conversely, the Neutral emotion model exhibits relatively weak explanatory power, suggesting that other unmeasured factors may play a larger role in determining Error in this condition. Across all models, RT consistently predicts Error, with faster responses associated with reduced precision, while N-Back has no significant impact.

Figure 4, demonstrates the varying influence of image number on Error across different emotional states. The significant relationship for Happy and Sad emotions highlights the importance of Intensity as a predictor of Error, while the weak relationship for Neutral emotion suggests that other factors may play a more substantial role. Overall, these models highlight the varying influence of RT, Intensity, and N-Back on Error across different emotional states, with Intensity and RT being more consistently significant predictors.

**FIGURE 4.**
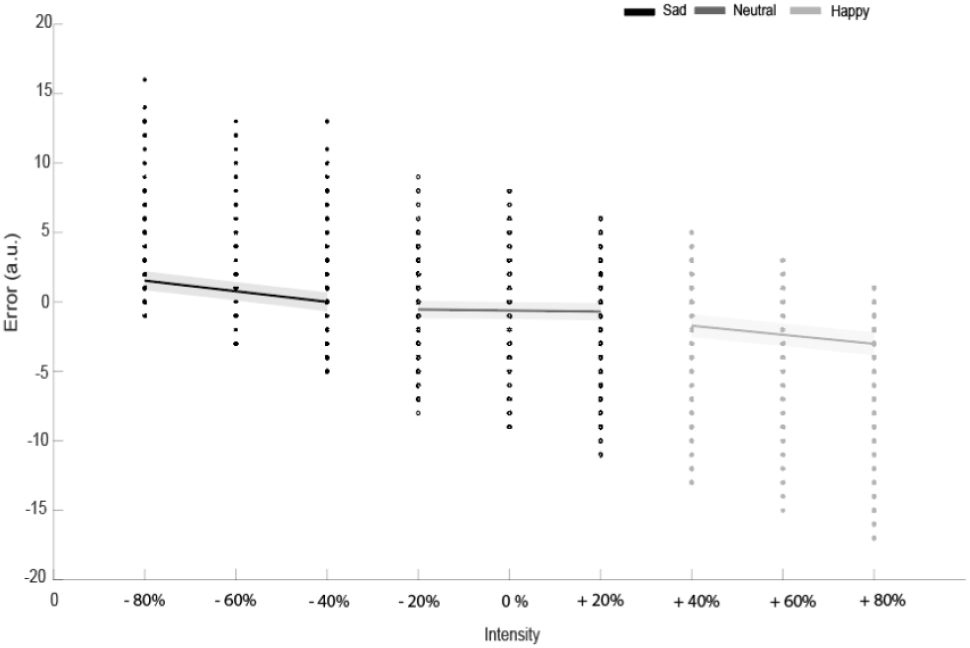
Linear Regression Fitted Errors by Emotion across valence axis. Scatter points, represent individual participant error at each intensity level. Fitted lines, show the predicted trend of error as number of image increases (image 1 = 90% sad or – 90%, image 19 = 90% happy or + 90%), based on linear regression. Shaded areas are confidence intervals, indicate the 95% confidence range around the fitted lines, reflecting the uncertainty in the regression predictions.

## 4. Discussion

Our study investigated how emotional valence and intensity, as well as cognitive load, influence perceptual accuracy and reaction times (RTs) in a delayed-reproduction task involving emotional morphed faces. The findings underscore the nuanced interplay between emotion and WM, contributing to our understanding of emotional biases and their implications for cognitive processing.

Our results revealed that emotional valence significantly influences perceptual accuracy. Participants exhibited higher error rates when recalling happy images compared to sad or neutral images. This finding aligns with evidence suggesting that happiness, despite being associated with positive emotional states, can reduce cognitive performance by broadening attention and increasing distractibility (Fredrickson, 2001; Rowe et al., 2007). Conversely, sadness, often linked to negative emotional states, appears to enhance focus, potentially reducing error rates by narrowing attention and fostering more detailed processing (Gasper & Clore, 2002). These findings support the notion that negative emotions such as sadness may aid cognitive performance in specific contexts, consistent with theories of negativity bias. Negative stimuli, including sad or angry faces, are often processed more efficiently due to their evolutionary relevance in threat detection and survival (Vaish et al., 2008; Öhman et al., 2001). However, the higher errors observed for happy faces suggest a “positivity penalty,” in which the lighter cognitive demands associated with positive stimuli may lead to less precise memory encoding or retrieval.

One intriguing finding was the valence reversal observed in participants’ recall of emotional faces. Happy images were recalled as less pleasant and sad images as less unpleasant than their initial presentation. This phenomenon may reflect the role of cognitive reappraisal, a strategy in which individuals reinterpret emotional stimuli to modify their emotional impact (Gross, 2015). Positive reappraisal, such as downplaying sadness, and negative reappraisal, such as emphasizing the drawbacks of happiness, could explain these shifts in perceived valence (Flores-Torres et al., 2022; Tsai et al., 2024). Valence reversals challenge conventional expectations of positive recall for positive stimuli and negative recall for negative stimuli. They underscore the complex influence of emotional context and cognitive processes on memory encoding and retrieval, suggesting that emotions are not static but dynamically reinterpreted based on situational factors (Wang & Yin, 2023).

Our findings provide further evidence for both negativity and positivity biases. The negativity bias, observed in faster and more accurate responses to sad faces, reflects the evolutionary importance of negative stimuli in threat detection and adaptive behavior (Fox et al., 2000; Vaish et al., 2008). Conversely, the positivity bias, supported by the happy-face advantage, highlights the preferential processing of positive stimuli due to their higher frequency and social relevance (Feyereisen et al., 1986; Öhman et al., 2001). These biases are not mutually exclusive and may operate simultaneously, influenced by task demands, stimulus properties, and individual differences. For instance, the negativity bias may dominate in tasks requiring rapid threat detection, while the positivity bias may emerge in contexts emphasizing social interactions or long-term memory (Adolphs, 2002; Mikels & Reuter-Lorenz, 2019).

As previously discussed, sad targets tend to be perceived as more pleasant, whereas happy targets are perceived as less pleasant as they are. This pattern is reflected in a shift in error patterns. This trend is further elucidated by the findings from the GLMM analysis. For higher intensities of sadness, such as 80% sad, the error rate increases compared to other intensity levels. This suggests that as the intensity of sadness increases, it is perceived as progressively more pleasant, potentially contributing to greater categorization errors. Conversely, for happy images, higher intensities, such as 80% happy, are associated with fewer errors, indicating that such stimuli are perceived as less pleasant than lower-intensity happy images. These findings suggest a perceptual asymmetry, where participants are less likely to select responses representing extreme levels of sadness or happiness, potentially due to the cognitive and affective demands of processing high-intensity emotions. Interestingly, the effect of intensity was less pronounced for neutral faces, which may reflect their lack of emotional salience. This aligns with prior research indicating that neutral faces are processed more variably, often influenced by the emotional context of preceding stimuli (Kauschke et al., 2019; Liberman, Manassi, & Whitney, 2018).

Reaction times (RTs) to emotional faces can vary based on the task and emotional context, reflecting differences in attentional focus and processing priorities. In our study, participants responded faster to sad faces compared to happy faces, which aligns with research suggesting that negative emotions like sadness may heighten vigilance, narrow attentional focus and facilitate processing speed, potentially reflecting an adaptive mechanism for managing negative emotions (Tipples, 2019). Conversely, the “happy-face advantage” is often observed in studies focusing on emotion recognition tasks, where happy faces are processed more quickly due to their visual salience and association with positive social cues (Esteves & Öhman, 1993; Srinivasan & Gupta, 2011). These conflicting patterns highlight how task-specific demands, such as attention to detail or broader contextual processing, influence emotional face perception.

Contrary to expectations, cognitive load, as manipulated through the N-Back conditions, did not significantly influence error rates or RTs. Although errors in the 2-back condition were descriptively higher across all emotional categories, these differences were not statistically significant, as confirmed by ANOVA. This aligns with findings by Holmer et al. (2024), who reported that mimicry interference primarily affects tasks with lower cognitive demands (e.g., 1-back). The trend observed in the 2-back condition may be attributable to primacy and recency effects, where items at the beginning or end of a sequence are more easily recalled (Sternberg & Sternberg, 2017; Glanzer & Cunitz, 1966). The lack of significant differences across N-Back levels may also reflect the relatively small sample size or the specific design of the WM task. Future research should explore whether increasing task difficulty or incorporating additional WM measures, such as complex span tasks, could better capture the interaction between cognitive load and emotional processing.

## Author contribution

All authors contributed to the article and approved the submitted version.

## Acknowledgments

We would like to gratitude to Ali Asgari and Mohsen Honar for their feedback on task design.

